# Haptoglobin and glutamine synthetase may biomark cachexia induced by anti-acute myeloid leukemia chemotherapy

**DOI:** 10.1101/2024.07.24.604689

**Authors:** Dean G. Campelj, Cara A. Timpani, Guinevere Spiesberger, Luke E. Formosa, Joel R. Steele, Haijian Zhang, Ralf B. Schittenhelm, Lewis Leow, Craig A. Goodman, Emma Rybalka

**Affiliations:** Institute for Health and Sport, Victoria University, Melbourne, Victoria, 8001, Australia; Biology of Ageing Laboratory, Centre for Healthy Ageing, Centenary Institute, Camperdown, New South Wales, 2050, Australia; Faculty of Medicine and Health, Charles Perkins Centre, University of Sydney, Sydney, 2050, Australia; Inherited and Acquired Myopathies Program, Australian Institute for Musculoskeletal Science, St Albans, Victoria, 3021, Australia; Department of Medicine—Western Health, Melbourne Medical School, The University of Melbourne, St Albans, Victoria, 3021, Australia; Department of Biochemistry and Molecular Biology, Monash Biomedicine Discovery Institute, Monash University, Clayton, Victoria, 3168, Australia; Monash Proteomics and Metabolomics Platform, Department of Biochemistry and Molecular Biology, Monash University, Clayton, Victoria, 3168, Australia; Centre for Muscle Research, and Department of Anatomy and Physiology, The University of Melbourne, Parkville, Victoria, 3010, Australia

**Keywords:** Anti-cancer chemotherapy, cachexia, skeletal muscle wasting, atrophy, biomarkers, haptoglobin

## Abstract

**Background:** Anti-cancer chemotherapy is an underappreciated contributor to cancer cachexia, an often irreversible body-wasting condition that causes 20-30% of cancer-related deaths. An obstacle to predicting, monitoring and understanding the mechanisms underlying chemotherapy cachexia is that each cancer (and sub-type) is assigned different chemotherapeutic compounds, typically in multi-agent regimens. Here, we investigate the chemotherapy induction regimen (CIR) used in the haematological cancer, acute myeloid leukemia (AML). We hypothesized that the AML CIR would induce cachexia, including loss of lean tissue mass and skeletal muscle atrophy.

**Methods:** Using an unbiased proteomics approach we interrogated the underlying molecular mechanisms. 3-month-old male Balb/c mice were treated with the AML CIR via intraperitoneal injections of daunorubicin (1.7 mg/kg) on days 1-3, and cytarabine (33.2 mg/kg) administered on days 1-7 or vehicle. Mice were assessed 24 hours after the last treatment, on day 8, or allowed to recover for 2 weeks and assessed on day 22. A third cohort was given access to running wheels in cages. We assessed body composition, whole body metabolism and assessed the muscle proteome using quantitative tandem mass tag labelling LC-MS/MS analysis.

**Results:** The AML CIR-induced acute cachexia involved a ∼10% loss of body mass, ∼10% loss of lean mass and ∼20% reduction in skeletal muscle fibre size. Whole body metabolism and ambulatory activity declined. This cachexic phenotype did not recover over the 2-week post-CIR period (lean mass loss post-CIR: 1 week ∼7% vs 2 weeks ∼9%). In voluntarily active CIR-treated mice, body wasting was exacerbated due to unchecked loss of fat mass (CIR sedentary: ∼31% vs CIR active: ∼51%). Muscle proteome studies revealed upregulation of haptoglobin (Hp) and glutamine synthetase (Glul), which were positively correlated with body and lean mass loss. Hp was sensitive to the conditional induction, recovery and exacerbation of AML CIR-mediated cachexia, suggestive of biomarker potential.

**Conclusions:** The AML CIR induces an acute reduction of body, lean and fat mass underpinned by skeletal muscle atrophy, hypermetabolism and catabolism. Our data uncovered a conditionally sensitive muscle biomarker in Hp, which may be useful as a prognostic tool across other scenarios of chemotherapy-induced myopathy and cachexia or as a target for therapeutic discovery in follow-up studies.

## Introduction

Cancer-associated cachexia is a multifactorial wasting syndrome of body and lean tissue mass that may also include fat loss [1] and anti-cancer chemotherapy has gathered traction as a critical cachexia-promoting factor [2]. Common hallmarks are skeletal muscle wasting, dysregulated metabolism and reduced food intake [3]. Patients with haematological cancers such as acute myeloid leukemia (AML) are prone to cachexia, although the phenomenon is less studied than with solid tumours [4]. Since skeletal muscle mass is a key prognostic factor for survival in AML, preventing muscle cachexia is critical for improving survival in a disease that already has grim survival statistics [4].

Cachexia in the AML setting may, in part, be driven by the intensity of initial treatment. Universally, this comprises the ‘7+3’ chemotherapy induction regimen (CIR) involving 7 days of cytarabine (an antimetabolite) concomitant with anthracycline (typically daunorubicin or idarubicin) on days 1-3 [5]. AML patients may receive multiple cycles of the CIR before complete remission is achieved. Thereafter, consolidation chemotherapy is administered and patients are assessed for haematopoietic cell transplantation (HCT), the only current curative strategy for AML. Pre-HCT cachexia leads to poor treatment-related outcomes and is a growing concern for clinicians [6]. Currently there are limited data on the impact of CIR on skeletal muscle (or on driving cachexia) and whether complete recovery from it is possible. Indeed, children with haematological cancers who receive intense chemotherapy tend not to recover their muscle mass and function, leaving survivors burdened with poor muscle health throughout life [7].

The inability to predict those likely to develop muscle cachexia and those who manifest early symptoms hampers the clinical treatment of cancer and other chronic diseases in which cachexia is devastating. In this regard, quantitative proteomics enables the detection of protein biomarkers that become disproportionate relative to the muscle-specific proteome in disease states. These proteins have the potential to not only indicate the likelihood and scope of cachexia impact, but also to predict the likelihood that cachexia could be life-threatening and assess the effectiveness of potential therapeutics. In this study, we sought to identify putative protein biomarkers of cachexia via quantitative tandem mass tag (TMT)-labelling proteomics. We hypothesised, in line with data for the pro-cachexia anthracycline analogue, doxorubicin [8], that the AML CIR would drive cachexia, including skeletal muscle wasting.

## Methods

### Experimental protocols and treatments

#### Animals

Three-month old (sexually mature) male Balb/c mice were acquired from the Animal Resource Centre (now Ozgene, Western Australia). Sex was controlled since AML is more prevalent in males. Mice were housed on a 12-hour light/dark cycle with *ad libitum* access to standard rodent chow and water.

#### Chemotherapy

Mice were randomly allocated to treatment groups: intraperitoneal injections of daunorubicin (1.7 mg/kg) on days 1-3 and cytarabine (33.2 mg/kg) on days 1-7, or delivery vehicle (VEH; 0.9% saline) daily (total *n*=20, 10/group). Daunorubicin dose was the maximum tolerable dose derived from our pilot studies based on dose escalation from a published starting dose [9]. Cytarabine dose was equivalent to clinical AML treatment [10] adjusted for mice based on FDA guidelines [11]. Mice were individually housed in metabolic cages to assess ambulatory and metabolic activity during CIR treatment. The experimental endpoint was 24 hours after the final chemotherapy treatment (i.e., day 8).

#### Recovery from chemotherapy

Communally housed mice (*n*=48, *n*=8/group, *n*=4-5/cage) received VEH or CIR treatment as stated above were assessed for recovery over time at 24 hours (day 8), 1 week (day 15) and 2 weeks (day 22) after the final CIR treatment (Fig 5A). There were no interventions during recovery.

#### Biomarker lability using exercise

VEH and CIR-treated mice were randomly allocated to individual metabolic cages containing a running wheel (total *n*=0, 10/group). ACT mice had *ad libitum* access to the running wheel throughout treatment (Fig 5A).

### Indirect calorimetry & activity monitoring

Promethion Metabolic cages fitted with laser tracking capacity and running wheels (Sable Systems, USA) were used to assess cage- and wheel-based physical activity and whole-body metabolism in real time, and to apply exercise. Mice acclimatised for 3 days and data collection occurred on days 4-10 across the 7-day treatment period [12].

### Body composition analysis

Echo Magnetic Resonance Imaging (echoMRI; EMR-150, Echo Medical Systems, USA) was used to assess body composition [12]. Live mice were scanned on day 1 (pre-treatment) and day 8 (post-treatment), and for recovery experiments, additionally on days 15 and 22 of the experiment.

### Tissue collection

24 hours after the last CIR treatment on day 7, and live analyses on day 8, mice were deeply anaesthetised with isoflurane (5% induction and 2-3% maintenance). Muscles and organs were surgically removed, weighed and snap frozen. Tibialis anterior (TA) muscles were prepared for histology.

### Histological analyses

TA muscles were cryopreserved in optimal cutting temperature compound (Sakura Finetek) using liquid nitrogen-cooled isopentane and cryosectioned at 8µm (−18°C, Leica CM1950). Mounted sections were stained with haematoxylin and eosin (H&E) or picrosirius red to assess fibre size and myopathy, and fibrosis, respectively. Slides were imaged on a Zeiss Axio Imager Z2 microscope (GmbH, Germany) at 20x magnification. ImageJ software (NIH, USA) was used for data acquisition as performed previously [12].

### Tandem Mass Tag (TMT)-labelled Proteomics and bioinformatics

TMT labelling proteomics on quadriceps samples were performed as previously described [13] as per Monash Proteomics and Metabolomics Platform data-dependent methodology [14]. Bioinformatics, Reactome (v83)-based pathways enrichment and deep pathways probing was performed as previously described [13]. Our analysis approach enables the detection of alterations in pathways that could remain unobserved when scoping only differentially expressed proteins because individual proteins might not provide comprehensive insights into how a specific pathway is changing when assessed through global, untargeted proteomic-level statistical comparisons. The FDR of <0.05 is reported.

### Statistics

Data are presented as mean ± standard error of the mean and were analysed with GraphPad Prism (v8, CA, USA; α=0.05). CIR versus VEH comparisons were analysed by t-test or two-way ANOVA (with repeated measures) as necessary. For CIR versus VEH recovery time course, data were analysed by repeated measures two-way ANOVA or mixed-effects analysis. For physical activity effects, two-way ANOVA (with repeated measures where necessary) was used. Tukey’s or Bonferroni’s post-hoc testing was applied to detect between group effects. Proteomics statistics are stated above. Linear regression correlations were performed on pooled sample sets to determine associations between biomarker proteins and anthropometric parameters.

## Results

### AML CIR induces acute cachexia in mice

AML CIR caused body mass decrements of 10%. Lean mass reduced by ∼10% and fat mass by ∼30% between pre- and post-treatment assessments (Fig. 1A). Body wasting was not entirely explained by reduced caloric intake since mean food intake in CIR mice was not statistically significant (Fig. 1B). Mass of the extensor digitorum longus (EDL), soleus and TA muscles was also reduced (Fig. 1C). However, when corrected for body mass, there was no effect of CIR (Supp Table 1), highlighting that muscle wasting is proportionate to the overall rate of body wasting. CIR also reduced the mass of all organs assessed (Fig. 1C). In summary, our data show that AML CIR induces clinically defined cachexia in mice.

**Figure 1:**
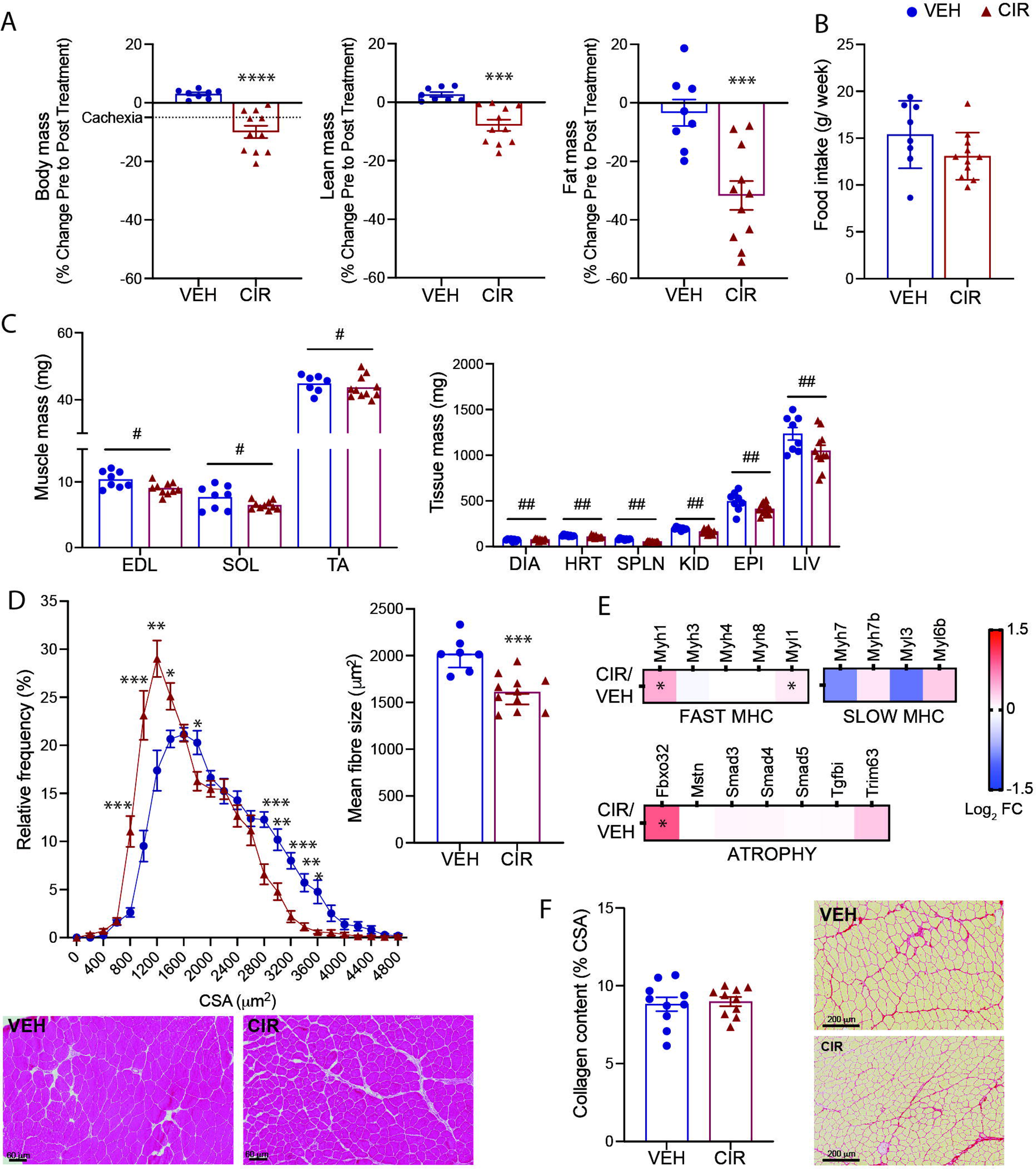
Clinically compatible cachexia profile of mice treated with anti-acute myeloid leukemia (AML) chemotherapy. Pre- to post-treatment changes in (A) body mass and composition of lean and fat mass. (B) Average daily food intake over the treatment protocol. (C) Effect of treatment on endpoint muscle and tissue mass and on (D) tibialis anterior (TA) fibre size distribution, mean fibre size and representative H&E-stained images. (E) Protein expression of fast and slow myosin heavy chain (MHC) isoforms and traditional atrophy signalling markers, Atrogin-1 (Fbxo32), TGFβ-myostatin-Smad axis, and Murf-1 (Trim63). (F) Collagen content and representative picrosirius stained images of TA. \**p*<0.05, \*\**p*<0.01, \*\*\**p*<0.001, \*\*\*\**p*<0.0001 CIR different from vehicle (VEH), #*p*<0.05, ##*p*<0.01 main CIR effect. Scale bar for H&E images = 50 µm; for picrosirius images = 200 µm.

**Table 1.** Response of muscle mass to voluntary exercise during acute myeloid leukemia (AML) chemotherapy induction regimen (CIR) Data are mean delta % change active (ACT) relative to sedentary (SED) ± SEM. EDL: extensor digitorum longus; SOL: soleus; TA: tibialis anterior. \**p*<0.05, \*\*\*\**p*<0.0001.

| | $\Delta$ % change VEH ACT v SED | $\Delta$ % change CIR ACT v SED |
| --- | --- | --- |
| EDL | $5.75 \pm 3.03$ | $40.41 \pm 6.54****$ |
| SOL | $14.99 \pm 3.12$ | $0.44 \pm 5.79^*$ |
| TA | $1.93 \pm 2.74$ | $-2.26 \pm 1.37$ |

Histological fibre sizing of TA muscles revealed CIR induced muscle atrophy (Fig. 1D), with more small fibres (800-1400 mm^2^) and fewer large fibres (2800-3600 mm^2^; Fig. 1D). Overall, CIR reduced the mean fibre CSA by ∼20% (Fig. 1D). Through probing our TMT-labelled proteomics data set (quadriceps muscle), we detected significant (based on FDR from pathways enrichment) upregulation of fast-twitch fibre-specific MyH isoforms, Myh1 and Myl, and of atrophy regulator, Atrogin-1, indicating that fast-type fibre transformations may attempt to compensate for global fibre atrophy (Fig. 1E). There was no effect of CIR on E3 ubiquitin-protein ligase, muscle RING-finger protein-1 (MuRF-1/Trim63) or components of the inducible transforming growth factor β (Tgfbi)/myostatin (Mstn)/suppressor of mothers against decapentaplegic (Smad) axis (Fig. 1E). There were no signs of active necrosis or regeneration that would indicate CIR-induced muscle damage (Fig. 1D) or of fibrotic myopathy (Fig. 1F).

### AML CIR reduces physical activity and systemic energy expenditure

Cage-based activity tracking revealed CIR mice reduced their physical activity levels from day 3 of treatment and maintained significantly lower levels for the duration of the treatment period compared to VEH mice (Fig. 2A). Interestingly, energy expenditure only decreased from day 4 of treatment, indicating a 24 h window where energy expenditure is increased relative to physical activity levels (Fig. 2B). However, lean tissue mass-corrected energy expenditure was not different between VEH and CIR-treated mice (Fig. 2C) indicating energy expenditure is relative to lean tissue mass and physical activity may reduce to conserve both. Since both fat and lean mass catabolism were implicated in CIR-induced body wasting (Fig. 1), we calculated the respiratory quotient (RQ; VCO_2_/O_2_), a marker of preferential substrate utilisation and shifts thereof. The RQ revealed a significant shift in fat relative to carbohydrate metabolism in CIR mice at day 7 (Fig. 2D).

**Figure 2:**
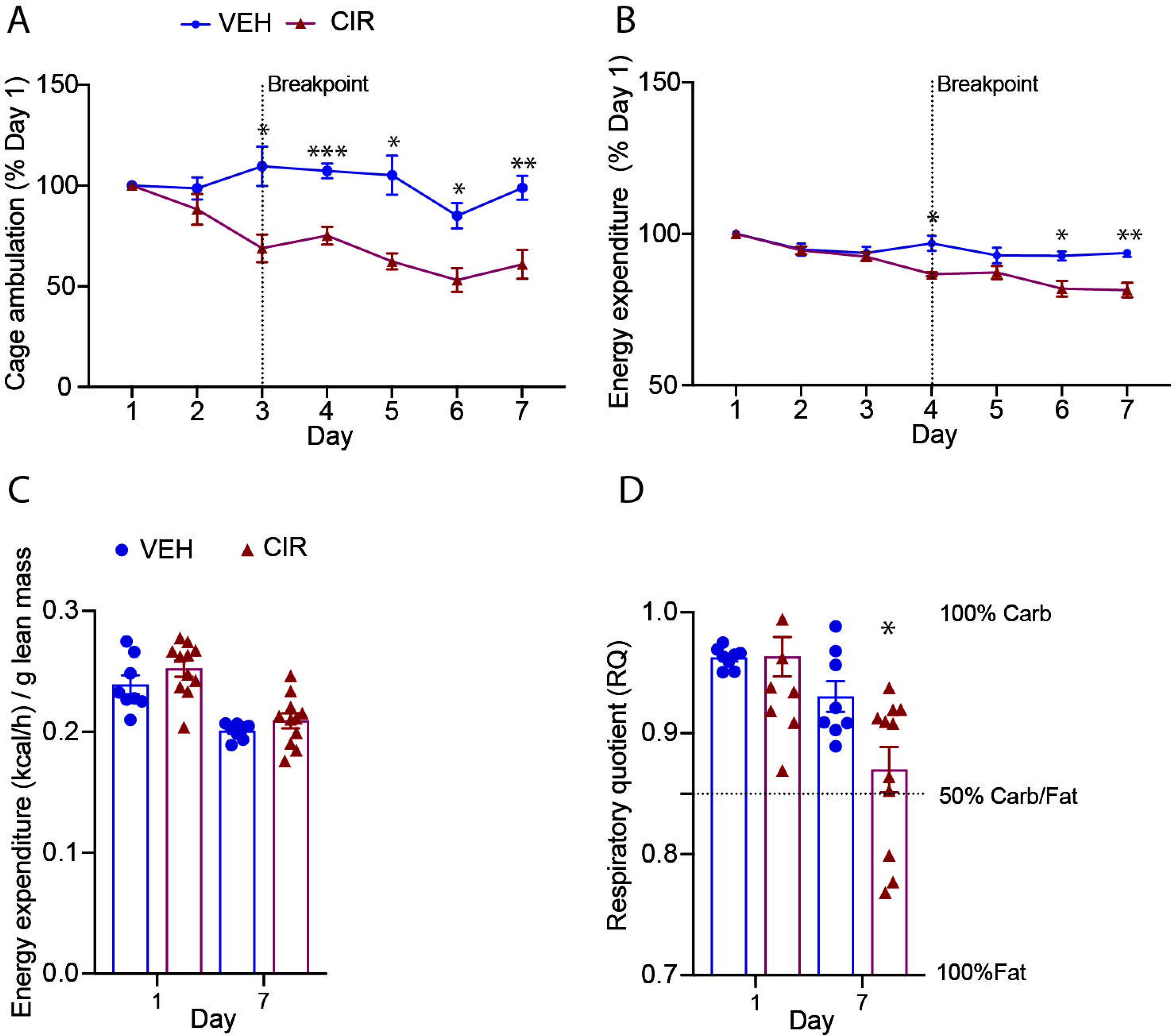
Physical activity and whole-body metabolism are reduced in mice treated with acute myeloid leukemia (AML) chemotherapy induction regimen (CIR) Mice were assessed continuously for (A) cage ambulatory distance and (B) energy expenditure, which is expressed relative to day 1 levels. (C) Start- and end-point energy expenditure (kcal) expressed relative to lean tissue mass and (D) the respiratory quotient (VCO_2_/VO_2_) are shown. Data are mean ± SEM. \**p*<0.005, \*\**p*<0.01, \*\*\**p*<0.001 CIR versus vehicle (VEH).

### Haptoglobin (Hp) and glutamine synthetase (Glul) are responsive to AML CIR-induced muscle cachexia

TMT-labelling proteomics enabled us to probe the molecular response underlying CIR-induced muscle wasting and to identify potential muscle-specific biomarkers. Of the 4,716 proteins detected, only Hp and Glul were differentially expressed in response to AML CIR treatment according to our cut-off criteria (log_2_ 0.75-fold change, adjusted *p* <0.05), and both were upregulated (Fig. 3A & B). Five additional proteins were different based on adjusted *p* (<0.05) only: Acad11 (acyl-coA dehydrogenase family member 11), Ca14 (carbonic anhydrase 14), Cd36 (platelet glycoprotein 4) and Plin4 (perilipin 4) were upregulated, while Dgkz (diacylglycerol kinase zeta) was downregulated. Pathways enrichment identified 12 significantly upregulated pathways in response to AML CIR (based on FDR; Fig. 3C) and all involved mitochondrial metabolism of amino acid, glycolytic and fatty acid substrates. The most significantly dysregulated pathway was branched chain amino acid (BCAA) catabolism, consistent with the loss of muscle mass observed. We deeply probed the BCAA, β-oxidation and mitochondrial tricarboxylic acid (TCA) cycle pathways to decipher mechanisms (Fig. 3D). Of the 18 proteins within the BCAA catabolism pathway our proteomics detected, CIR upregulated 14 (78%). The most significantly upregulated protein was branched chain keto acid dehydrogenase E1 subunit a (Bckdha), an inner mitochondrial protein involved in catabolism of leucine, isoleucine and valine. In fact, most of the 14 upregulated BCAA proteins were linked to catabolism of valine or leucine. There were 47 oxidation-related proteins captured in our proteome and 27 were upregulated (57%) by AML CIR treatment. Notably, almost all family members of acyl-coA dehydrogenase (Acad), an enzyme involved in the metabolism of acyl-coA variants, were upregulated (as was Acad6A1 in the BCAA pathways). Acad11, a recently characterised 4-hydroxy acid (4HA) acyl coA dehydrogenase that localises to peroxisomes and plays an important role in facilitating longer-chain 4-HA catabolism [15], was the most significantly upregulated protein within the β-oxidation pathway. Several protein subunits of the mitochondrial TCA cycle were upregulated (aconitase (Aco2), isocitrate dehydrogenase (Idh1) and malate dehydrogenase (Mdh2)), although notably the TCA pacesetter and mitobiogenesis biomarker, citrate synthase (Cs), was not. Our data suggest a remodelling of mitochondria and peroxisomal metabolism to drive BCAA (especially valine and iso-/leucine) and fat oxidation. Since anthracyclines are metabolised by mitochondrial Complex I (mit-CI) [16, 17], we also probed this pathway. Specific components of NADH-ubiquinone oxidoreductase (Nduf) were upregulated, including core subunit V2 (Ndufv2), which is known to protect against doxorubicin-induced mitochondrial dysfunction driven cardiomyopathy [18]. Uncoupling-related proteins were also probed as a potential mechanism of hypermetabolism (Fig. 3E). Uncoupling proteins 1 and 3 (Ucp1, 3) were both upregulated by CIR and Ucp1 more so than Ucp3 indicating thermogenesis as a mechanism.

One hundred and thirty-five Reactome pathways were downregulated in CIR versus VEH muscle (Supp Table 2) and the top 25 pathways are presented in Fig. 3F. Many of the downregulated pathways were associated with ribosomal protein translation and synthesis and the topmost downregulated pathway (based on FDR from pathways enrichment) was inflammation-dependent (via tumour necrosis factor receptor 1 (Tnfr1)) ceramide production. In fact, several pathways concerning inflammation regulator, nuclear factor kappa B (Nf-κb) activity were reduced. Of note, the macroautophagy pathway, which when conditionally up- or down-regulated can induce muscle atrophy [19], was significantly downregulated. Collectively, our data indicate that muscle protein synthesis pathways are suppressed, and catabolism is increased to provide substrates for mitochondrial metabolism.

**Figure 3:**
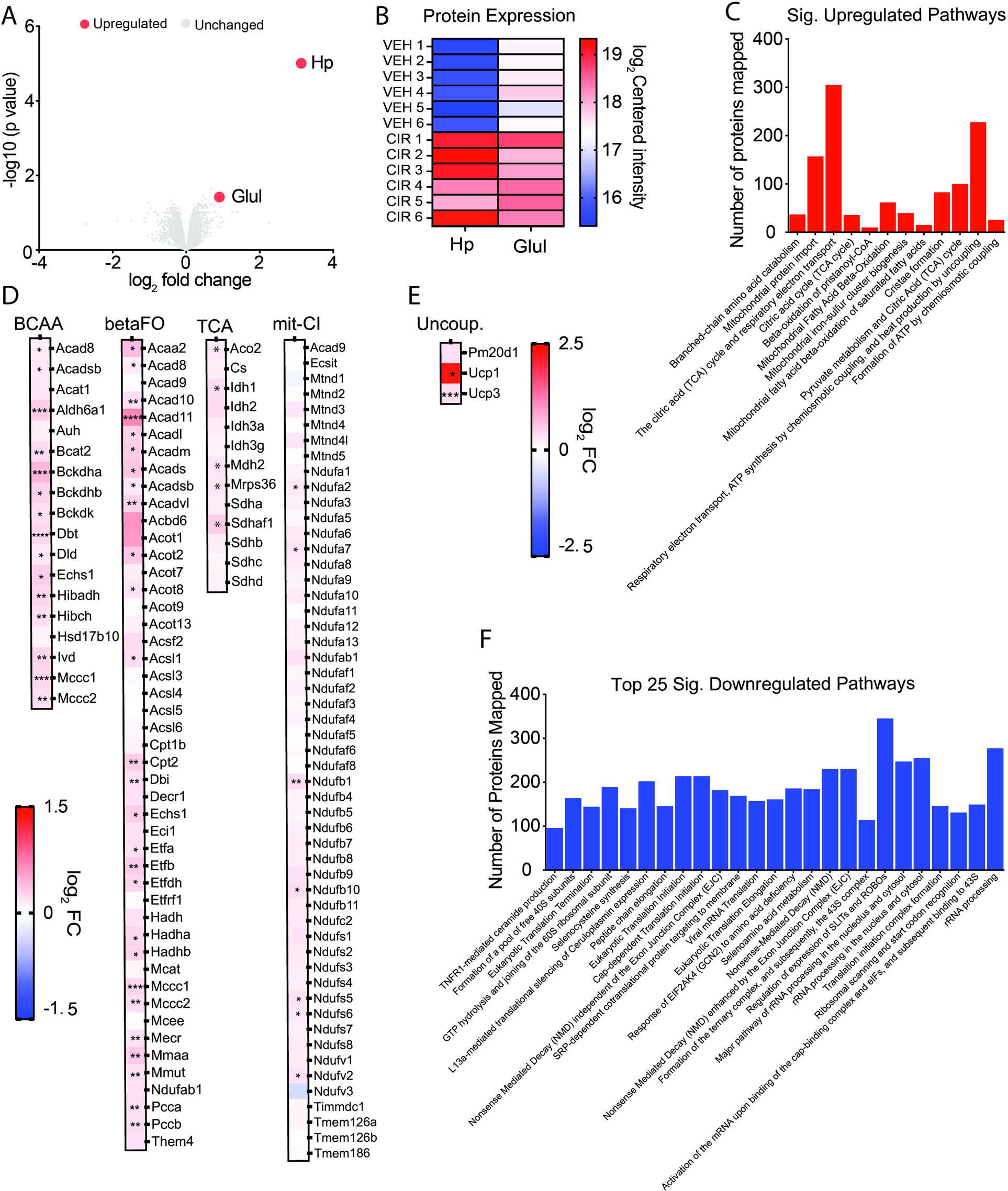
Muscle proteome response to acute myeloid leukemia (AML) chemotherapy induction regimen (CIR) reveals haptoglobin (Hp) and glutamine synthetase (Glul) as potential biomarkers. (A) Volcano plot showing upregulation of Hp and Glul based on 0.75 log_2_-fold change and adjusted *p* <0.05 cut-off. (B) Heatmap of individual mouse muscle expression of Hp and Glul. (C) Significantly upregulated Reactome pathways. (D) Protein expression profile of branched chain amino acid (BCAA) catabolism, and β-fat oxidation, mitochondrial tricarboxylic acid (TCA) cycle and mitochondrial Complex I (mit-CI) metabolism Reactome pathways. (E) Uncoupling-related protein expression. (F) top 25 downregulated Reactome pathways in response to AML CIR treatment. \**p*<0.05, \*\**p*<0.01, \*\*\**p*<0.001, \*\*\*\**p*<0.0001 CIR versus vehicle VEH based on *p* <0.05.

### Fat, but not lean mass, recovers in the two weeks following AML CIR

To determine whether body, lean and fat mass could recover after chemotherapy treatment, we next treated a cohort of mice with AML CIR then allowed 2 weeks of untreated recovery during which body mass and composition were assessed daily and weekly, respectively (Fig. 4A). Body mass was lowest at day 9 (2 days into recovery; ∼80% of starting body mass) and began to recover from day 10 (Fig. 4B). However, by the end of the 2-week recovery period, body mass had not fully recovered remaining at 90-95% of the starting body mass. Body mass displacement from the VEH group was greatest on day 8 and least on day 22 of recovery, however there was no significant difference between assessment time points in the CIR group (Fig. 4C). Body mass did not recover sufficiently to renounce the clinical definition of cachexia. Lean mass was more resistant to recovery than fat mass. While fat mass displacement from VEH was still statistically different at day 22 of recovery, it did partially recover between day 8 and 22 by ∼50%, whereas lean mass displacement did not shift (Fig. 4C). There was no recovery of EDL, soleus or TA muscle mass observed (Fig. 4D). The muscle proteome was also resistant. There was no significant change in muscle Hp or Glul expression at day 22 based on log-fold change and adjusted *p* value post-recovery (Fig. 4E). However, there were shifts in quadriceps Hp expression for 3/5 animals indicating Hp is responsive to withdrawal of CIR and may biomark early changes in molecular signalling within muscle (Fig. 4E). In contrast, Glul was non-responsive to CIR withdrawal (Fig. 4E).

**Figure 4:**
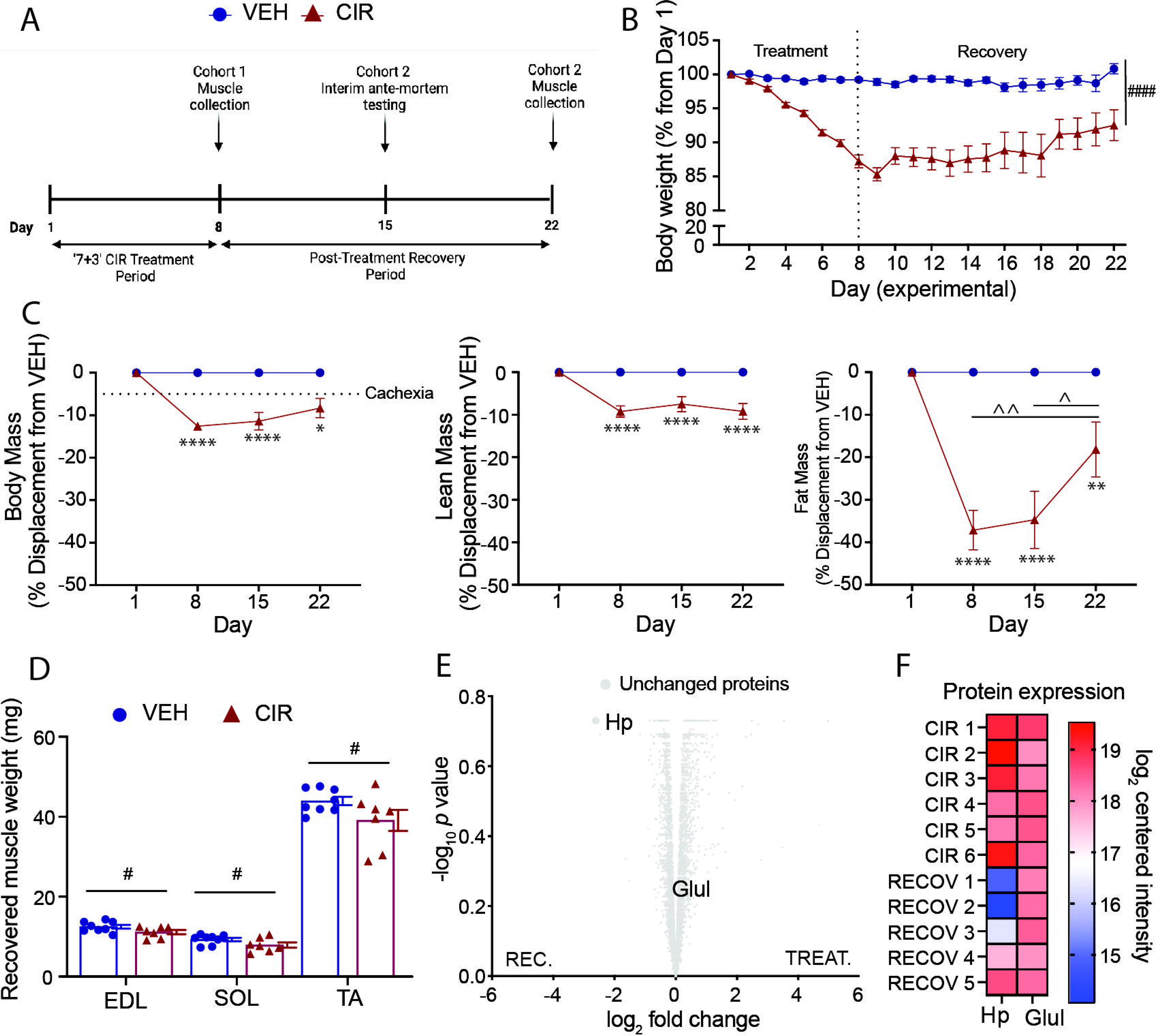
Recovery profile following of acute myeloid leukemia (AML) chemotherapy induction regimen (CIR) (A) Experimental timeline. (B) Body mass over the AML CIR and recovery time course. (C) Body, lean and fat mass displacement relative to vehicle (VEH) over the recovery time course. (D) Endpoint muscle mass of the extensor digitorum longus (EDL), soleus (SOL) and tibialis anterior (TA). (E) Volcano plot of unresponsive quadriceps muscle proteome to CIR cessation after two weeks based on log_2_-fold change and adjusted *p* <0.05 cut-off. (F) Haptoglobin (Hp) and glutamine synthetase (Glul) expression at the treatment (CIR) and recovery (CIR RECOV) endpoints for individual animals. \**p*<0.05, \*\**p*<0.01, \*\*\*\**p*<0.0001 CIR RECOV relative to age-matched VEH; #*p*<0.05, ####*p*<0.0001 main group effect; ^*p*<0.05, ^^*p*<0.01 within group time effect.

### Voluntary exercise during AML CIR treatment worsens cachexia and further upregulates Hp

We used *ad libitum* voluntary running activity (ACT) over the 7-day AML CIR to test (1) potential consequences during CIR and (2) the lability of Hp and Glul as cachexia biomarkers. On average, VEH ACT mice covered ∼100x more distance on the running wheel compared to ground metres covered by VEH SED mice, while CIR ACT mice covered ∼60x more than CIR SED mice (data not shown). From day 3 of treatment, CIR ACT mice became steadily less active (ground and wheel metres) with endpoint physical activity <50% of starting activity levels relative to VEH ACT (Fig. 5A). Energy expenditure reduced proportionate to ambulatory distance, although notably, a day earlier than physical activity decline (Fig. 5B). In healthy VEH mice, food intake increased to match the additional energy expenditure from increased physical activity (Fig. 5C) sufficient to resist changes in body mass (Fig 5D). In contrast, CIR ACT mice did not increase food consumption to meet energy expenditure (Fig. 5B and C). While incremental changes in body and lean mass were too small to be statistically different, CIR ACT mice trended to lose body and lean tissue mass (significant CIR group effects), whereas VEH ACT mice tended to reduce body mass but increase lean mass (Fig. 5D). The most significant effect of increased physical activity was on fat mass. VEH ACT mice lost ∼30% of their pre-ACT fat mass, whereas CIR ACT mice lost ∼50% for less activity. At the muscle level, the fast-twitch glycolytic type 2 EDL muscle of CIR ACT mice increased in size – the predominantly slow-twitch oxidative type 1 soleus did not change (Table 1). In contrast, in WT ACT mice, the EDL mass did not shift but the soleus mass increased (Table 1). There was no effect of exercise on mass (Table 1) or mean fibre size (Fig. 5E) of the mixed fibre type TA muscle in either VEH or CIR treated mice.

**Figure 5:**
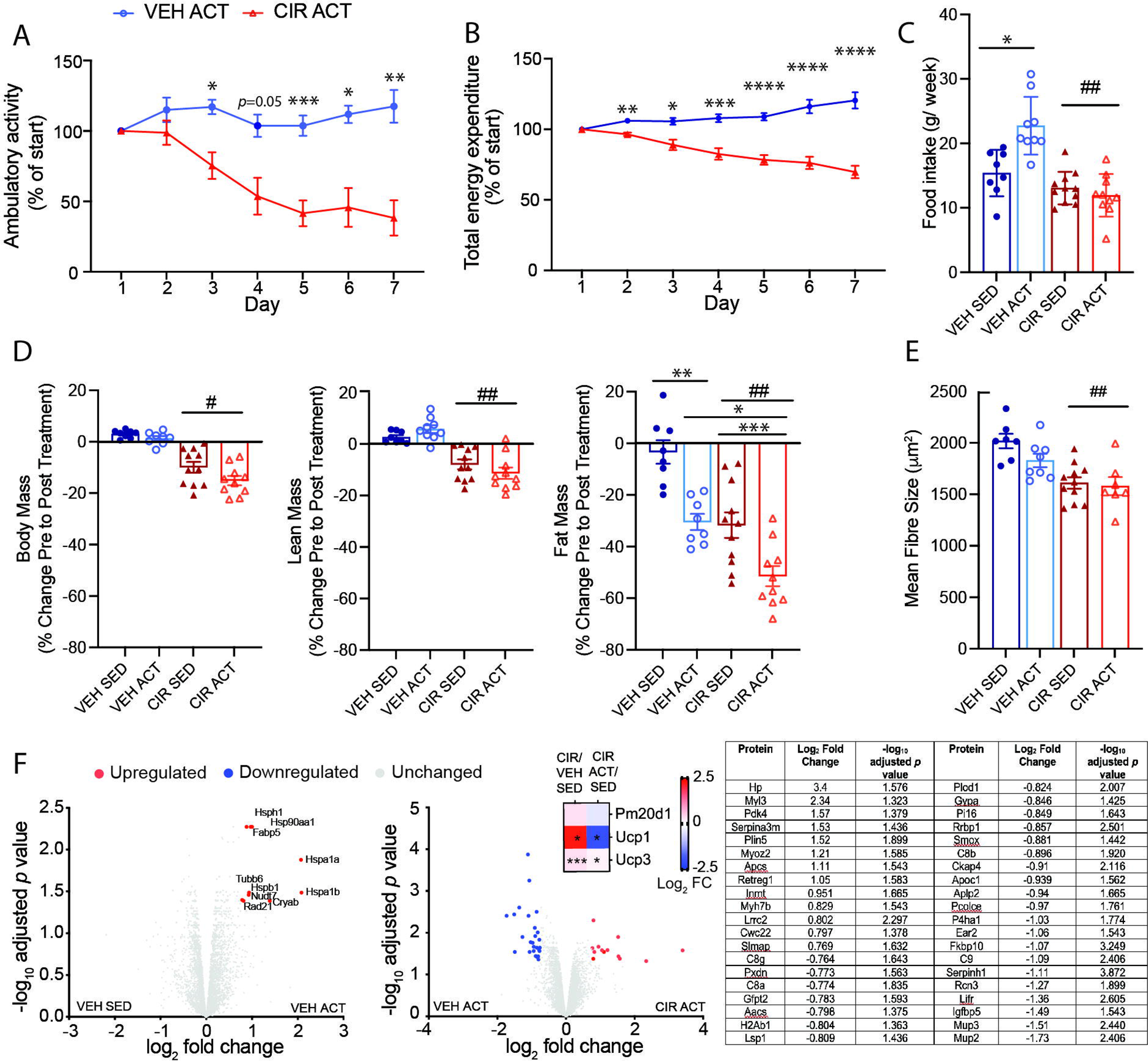
Worsening of cachexic fat mass loss by voluntary exercise during acute myeloid leukemia (AML) chemotherapy induction regimen (CIR) treatment further upregulates muscle haptoglobin (Hp) expression. (A) Voluntary physical activity (wheel running) reduces over the AML CIR treatment period alongside (B) reduced total energy expenditure. (C) Daily food intake and (D) change in body, lean and fat mass over the treatment period. (E) Mean tibialis anterior fibre size and (F) quadriceps proteome response to voluntary exercise in vehicle (VEH) and CIR treated mice based on log_2_-fold change and adjusted *p* <0.05, including reduction of uncoupling protein (Ucp) expression based on FDR <0.05 from pathways analysis. \**p*<0.05, \*\**p*<0.01, \*\*\**p*<0.001 active (ACT) versus sedentary (SED) and CIR versus VEH ACT; #*p*<0.05, ##*p*<0.01 main treatment effect.

Physical activity induced a unique proteomic signature (10/4,766 differentially regulated proteins) in VEH mouse quadriceps involving upregulation of heat shock, DNA repair, muscle regeneration, and fatty acid metabolism proteins (Fig. 5F). Relative to VEH ACT, the CIR ACT proteome was considerably more impacted with 40 differentially regulated proteins detected (13 upregulated, 27 downregulated). The most upregulated protein was Hp, which was 6.8-fold higher than VEH following exercise exposure (i.e., CIR ACT v VEH ACT) compared with 5.8-fold higher in sedentary mice (i.e., CIR SED v VEH SED). Other upregulated proteins were associated with slow fibre isoform, extracellular matrix remodelling, muscle regeneration, and energy metabolism. Downregulated proteins were associated with immunogenicity/inflammation, endoplasmic reticulum proliferation, collagen biosynthesis, and polyamine, fatty acid and cholesterol biosynthesis indicative of a whole cell wasting phenotype. Ucp1 expression was normalised and Ucp3 expression was downregulated by exercise. Muscle Glul expression was not different in the AML CIR ACT relative to the VEH ACT group.

### Interrogating Hp and Glul as biomarkers of cachexia induced by AML CIR

Hp was the most responsive and labile muscle protein to AML CIR treatment of the >4,700 detected within our proteome (Fig. 6A). Hp’s scaled heatmap signature for all conditions expressed relative to VEH SED best reflected the scope of cachexia seen across our experimental conditions, where expression was highest under conditions that caused the largest mean reduction in body, lean, fat and muscle mass (Fig. 6A). However, only moderate-weak correlations were observed between Hp and each of body and lean mass loss from pre- to post-treatment and there was no correlation with fat mass loss (Fig. 6B). Hp better correlated with lean mass (r^2^=0.458, *p*=0.0004; red points and regression line) when the recovery readouts were removed (denoted by red dots and regression line) indicating it may be a better predictor of CIR-specific molecular pathology and associated muscle mass decrements. There was a weak correlation between Hp expression and endpoint raw quadriceps mass (matched for the same muscle used for proteomics) when recovery data points were removed. Glul expression was less responsive to treatment than Hp, failing to respond to CIR withdrawal during the recovery period or the added stress of exercise on top of CIR. Glul expression increased in VEH quadriceps over the 2-week recovery period and due to physical activity, consistent with growth- and physical activity-related metabolism increases. Glul was strongly correlated with body mass (when recovery data were removed, moderately correlated with all data included), moderately correlated with lean mass, and weakly correlated with fat mass change from pre-post treatment indicating its expression is relative to the overall metabolic state based on body mass. Glu expression did not correlate with endpoint muscle mass. CIR-induced heatmap signatures of atrophy mediators, Fbxo32 (Atrogin-1) and Trim63 (Murf-1), mimicked the expression pattern of Hp but not Glul, suggesting Hp sensitively recapitulates atrophy signalling.

**Figure 6:**
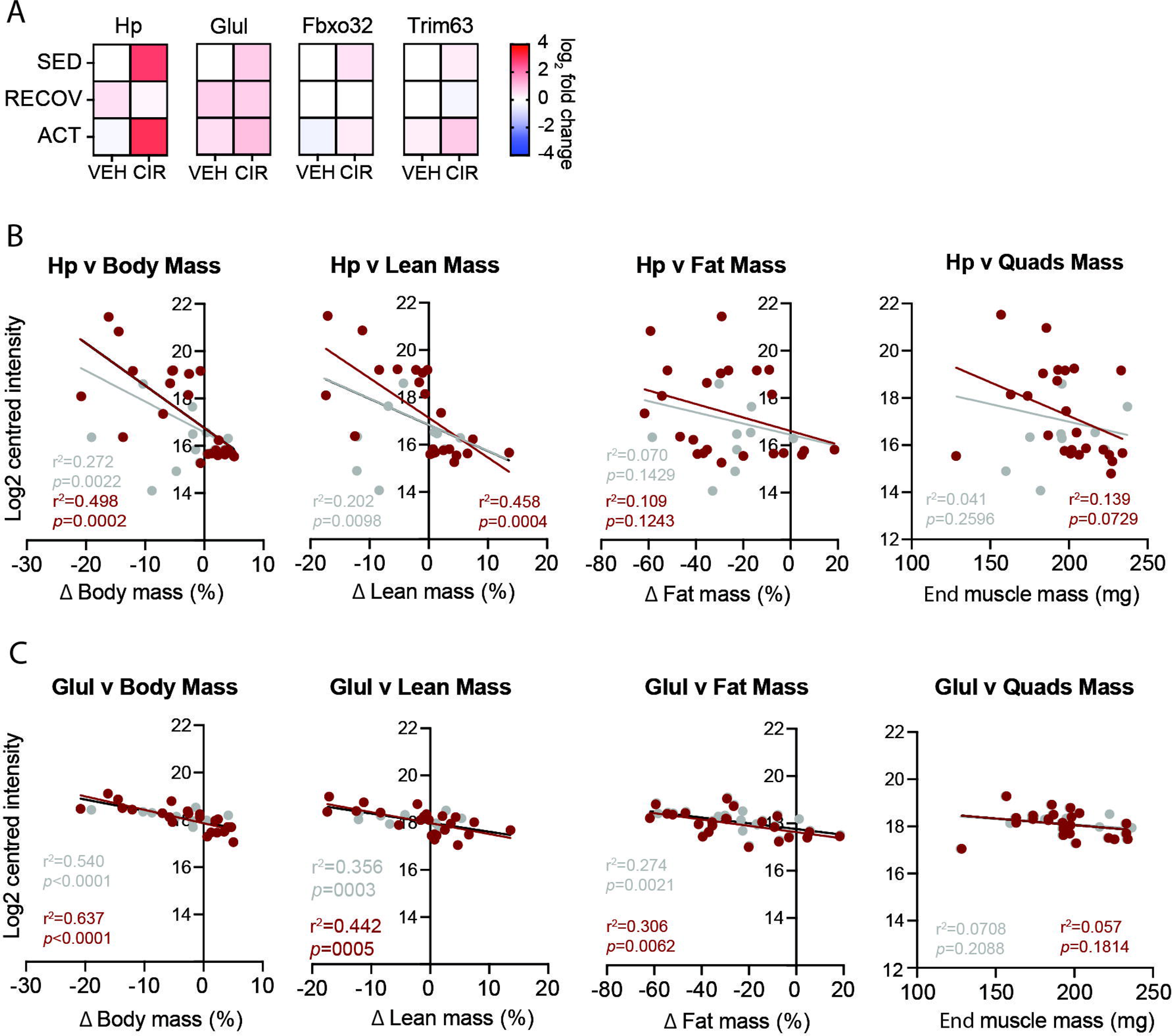
Biomarker potential of muscle haptoglobin (Hp) and glutamine synthetase (Glul) in the context of acute myeloid leukemia (AML) chemotherapy induction regimen (CIR) cachexia. (A) Conditional heatmap signatures of Hp, Glul, Fbxo32 (Atrogin-1) and Trim63 (Murf-1) in AML CIRL and vehicle (VEH) treated muscles. Correlations of (B) Hp and (C) Glul expression with each other and with change in body, lean, fat mass. Grey dots/regression lines denote data points from VEH and CIR groups under all conditions, red data points/regression lines denote VEH and CIR recovery data points removed.

## Discussion

Haematological cancers, such as AML, are rarely studied in the context of cachexia due to the rapid turnaround from diagnosis to initiation of chemotherapy treatment and evaluation for HCT and our study attempted to tackle this aspect. Despite only lasting one week, we reveal that the AML CIR induces a remarkable level of cachexia involving all the definitive hallmarks [1]. We show an acute reduction of body, lean and fat mass underpinned by skeletal muscle atrophy, hypermetabolism and muscle and fat catabolism. Mice appear to self-execute a tissue preservation mechanism by lowering physical activity and energy expenditure to reduce the catabolic insult akin to the human “sickness response” [4]. Persistent hypercatabolism of fat and muscle despite reduced activity and energy expenditure was a curious finding in our study but mechanistic clues emerged from our proteomics screen. From a fat mass perspective, Acad 11 and 10 – recently linked to coordinated lipid peroxidation-mediated 4-HA metabolism between peroxisomes and mitochondria [15] – were upregulated in our β-oxidation pathways analysis (Acad11 especially so). Lipid peroxidation is a well-established outcome of anthracycline-mediated reactive oxygen species production [17] and 4-HA species may be a byproduct, although it is unclear whether increased 4-HA metabolism is a defensive mechanism that spares muscle from lipotoxic myopathy, is purposeful to fuel thermogenic mitochondrial uncoupling, or both. These enzymes may be targetable to mitigate cachexia-related fat loss.

At the muscle level, upregulation of Glul, which enzymatically removes ammonia from the muscle purine nucleotide cycle during conversion of glutamate to glutamine [20], was a distinct outcome of CIR treatment. Glutamine release from skeletal muscle escalates during proteolysis [21] but cytarabine metabolism also generates significant ammonia via deamination [22] that may necessitate more Glul as an adaptive stress response. As muscle mass declines, cytarabine-generated ammonia load may overwhelm adaptive Glul overexpression and either drive, or be symptomatic of, muscle wasting. Ammonia accumulation is linked to multiple muscle wasting conditions [23]. Additionally, anthracyclines are metabolised by mit-CI and we speculated that subunit proteins might be upregulated to effectively process the daunorubicin load. NADH-ubiquinone oxidoreductase sub-units, especially subunit V2 (Ndufv2), were upregulated as previously shown in the context of doxorubicin treatment to protect against cardiomyopathy [18]. We also showed upregulation of Ucp 1 and 3, which drive heat over energy (and CO_2_) production explaining why catabolism increased while energy expenditure (measured via respirometry) reduced. It is difficult to differentiate whether mitochondrial uncoupling is fundamentally important to limit oxidative stress linked to daunorubicin metabolism by mit-CI or a thermogenic mechanism is activated as fat mass reduces and physical activity levels decline. In the latter scenario, mitochondrial uncoupling likely offsets an increased propensity for hypothermia. Our data suggest that a consequence of these chemotherapy-induced adaptations is inadvertent systemic hypermetabolic muscle catabolism. Defending the skeletal muscle BCAA pool from chemotherapy-induced ubiquitylation of L-Type amino acid transporter 1 (LAT1) is a promising approach recently explored *in vitro* [24] and may be useful in the context of the AML ‘7+3’ CIR. This mechanism may link back to upregulated Glul, whereby a futile increase in skeletal muscle glutamine exchange for BCAAs is facilitated by LAT1.

Through unbiased proteomic profiling of quadriceps muscle, we reveal two potential novel biomarkers of AML CIR-induced cachexia, Hp positively correlates with loss of body, lean and muscle mass and is highly responsive to conditional induction, recovery and exacerbation of AML CIR-mediated cachexia. Under normal physiological conditions, Hp is synthesised by the liver in response to erythrocyte degradation and circulated in the plasma as an acute-phase protein to detoxify free haemoglobin [25]. Hp’s role in skeletal muscle is poorly understood. Data from Zip14 metal transporter ablated mice suggest that muscle level expression is sensitive to inflammation-induced stress response signalling [26] and our data confirm Hp expression is highest when heat-shock proteins are also upregulated, i.e., with voluntary wheel running. In humans, plasma Hp is higher in atrophy resistant subjects exposed to 10 days of unloading and lower in cachexic cancer patients [27]. However, Massart *et al.* demonstrated increased skeletal muscle Hp expression in the C26-adenocarcinoma induced cachexia model [28] consistent with our data. Interestingly, Hp abundance is unaffected by Atrogin-1- and Murf-1-dependent immobilisation-atrophy suggesting that its transcription is invoked by a specific insult to skeletal muscle homeostasis [29]. Oxidative stress-associated protein carbonylation is apparent in Hp^-/-^ KO mice with muscle atrophy, weakness and fatigue, indicating Hp is redox sensitive [30]. Our data highlight that muscle specific Hp expression patterns mimic mitochondrial stress levels and iron accumulation, a hallmark of doxorubicin-induced cardiotoxicity [31], i.e., our pathways analysis revealed upregulated iron sulfur cluster biogenesis. That Hp levels were reducing in some mice by the end of the post-CIR recovery period indicates its sensitivity to CIR effects on muscle. In contrast, Glul was positively correlated with body wasting but was unresponsive to conditional recovery or exacerbation of CIR-induced cachexia by exercise. It appears to biomark overall metabolism, but not CIR-induced hypercatabolism specifically. Further evaluation of these proteins as predictive and/or prognostic biomarkers of cachexia involving muscle wasting are warranted. While a useful biomarker would typically detect in biofluids (i.e., blood, urine), there is opportunity for muscle specific biomarkers to be clinically useful across cancer treatment since AML treatment involves minor surgical procedures that enable access to muscle. We cannot rule out that muscle Hp is secreted into the bloodstream which would make it a particularly useful biofluid marker of cachexia. There is no evidence that Glul is secreted from muscle, however, plasma glutamate/glutamine ratios may predict muscle Glul activity and the cachexia hypercatabolic state.

It was surprising that exercise exacerbated cachexia by driving fat mass loss without beneficial recovery of lean tissue in our CIR treated mice. We saw shifts in muscle mass in fast-twitch but not slow-twitch muscles with CIR treatment, which directly opposed exercise effects observed in VEH mice, indicating that mitochondria-dense slow twitch fibres may be predominantly affected by CIR combined with the specific muscle activation pattern used in wheel running. Exercise, did, however, reduce Ucp 1 and 3 expression suggesting that over the longer term, it may be useful to abate hypermetabolism if mitochondrial uncoupling is the primary cause. Exercise is generally positively associated with muscle mass, function and metabolism and is being explored as a strategic approach to prevent cachexia [32]. Our data indicate the modality and potentially, the intensity, of exercise selected for AML patients while they are being actively treated is an important consideration. Previously, Wehrle *et al*. compared endurance and resistance training application during chemotherapy induction in 22 (of 29; 24% dropout) acute leukemia patients and demonstrated that only resistance training improved knee extension strength whereas endurance and no training reduced it [33]. The study did not measure body composition nor report on whether any participants were, or became, cachexic. In breast cancer patients undergoing chemotherapy treatment, 12 weeks of aerobic exercise (walking) reduced body and fat mass like our study, although notably, cachexia is not so much a problem in this malignancy or with its treatment [34].

## Conclusions

Importantly, we demonstrate the cachexic phenotype of an AML CIR cachexia mouse model illuminating hypermetabolism of BCAAs and fat as drivers through metabolic rewiring and mitochondrial uncoupling. We identified Hp and Glul as potential muscle specific biomarkers that could be developed as prognostic tools to monitor efficacy of interventional therapeutics against cachexia. Validation studies are necessary to map Hp and Glul expression against cachexia and recovery time course. Our two-week recovery period was insufficient in this regard. Future studies characterising the CIR-induced phenotype would benefit from: 1) body mass recovery-directed endpoints; 2) investigation of multiple CIR impact as is often used clinically; and 3) daily muscle and blood sampling across the AML CIR to tease out the contributions of each of daunorubicin and cytarabine to muscle proteome changes. Our data also illuminate potential therapeutic avenues against hypermetabolism in the metabolic pathways surrounding Hp (iron metabolism), Glul (urea processing and glutamate/glutamine metabolism), Acad 10 and 11 (4-HA metabolism) and mitochondrial uncoupling for follow-up.

## Acknowledgements

Grant support from the Institute for Health and Sport, Victoria University was awarded to DGC. This study used Bioplatforms Australia-/ National Collaborative Research Infrastructure Strategy-enabled infrastructure located at the Monash Proteomics and Metabolomics Platform.

## Ethical standards

Animal studies were approved by the Victoria University Animal Ethics Committee (AEETH17/017) and conformed to Australian standards.

## Supplemental tables

**Supplemental Table 1:** Muscle and organ mass/body mass (mg/g) ratios. Data are mean ± SEM.

|  | VEH | CIR |
| --- | --- | --- |
| *Muscles* |  |  |
| Extensor Digitorum Longus<br>(EDL) | 0.419 ± 0.007 | 0.451 ± 0.011 |
| Soleus (SOL) | 0.299 ± 0.013 | 0.307 ± 0.010 |
| Tibialis Anterior (TA) | 1.787 ± 0.029 | 1.903 ± 0.025 |
| *Tissues/Organs* |  |  |
| Diaphragm (DIA) | 2.834 ± 0.151 | 3.421 ± 0.185 |
| Heart (HRT) | 4.718 ± 0.128 | 4.742 ± 0.103 |
| Spleen (SPLN) | 3.206 ± 0.118 | 2.421 ± 0.059 |
| Kidney (KID) | 7.655 ± 0.149 | 7.201 ± 0.165 |
| Epididymal Fat (EPI) | 19.340 ± 1.466 | 18.065 ± 0.624 |
| Liver (LIV) | 48.088 ± 2.057 | 45.781 ± 1.630 |

**Supplemental Table 2:**
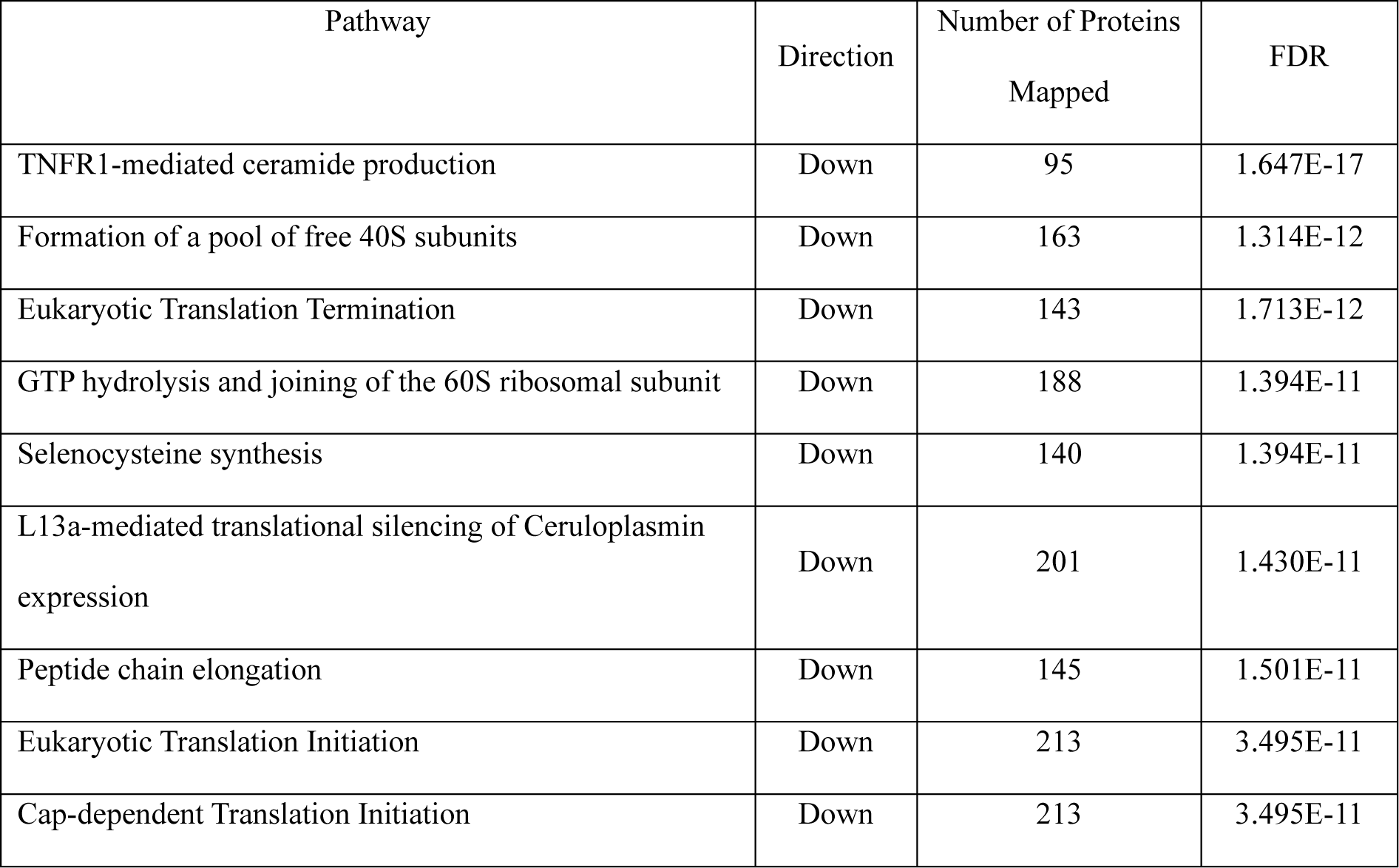
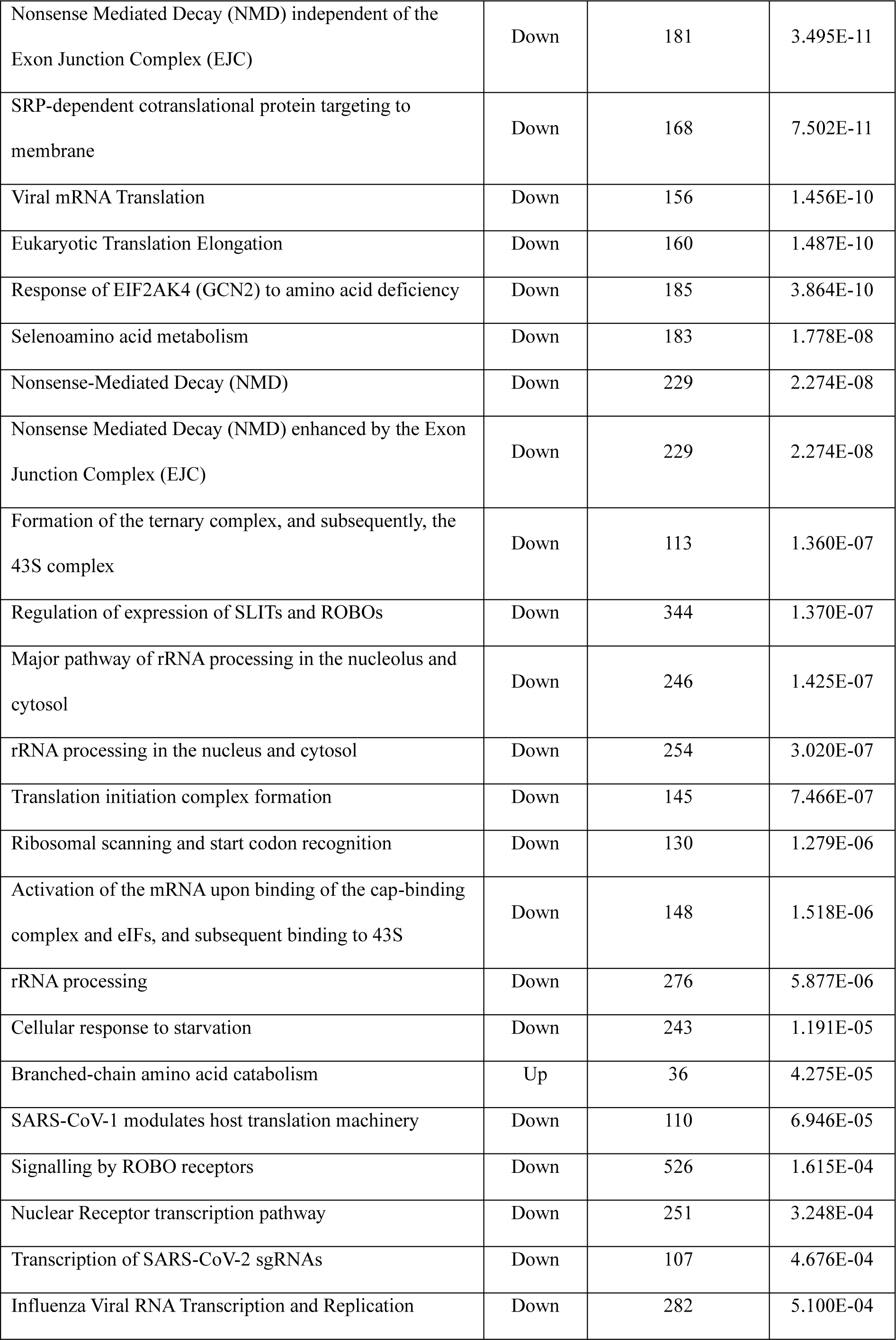
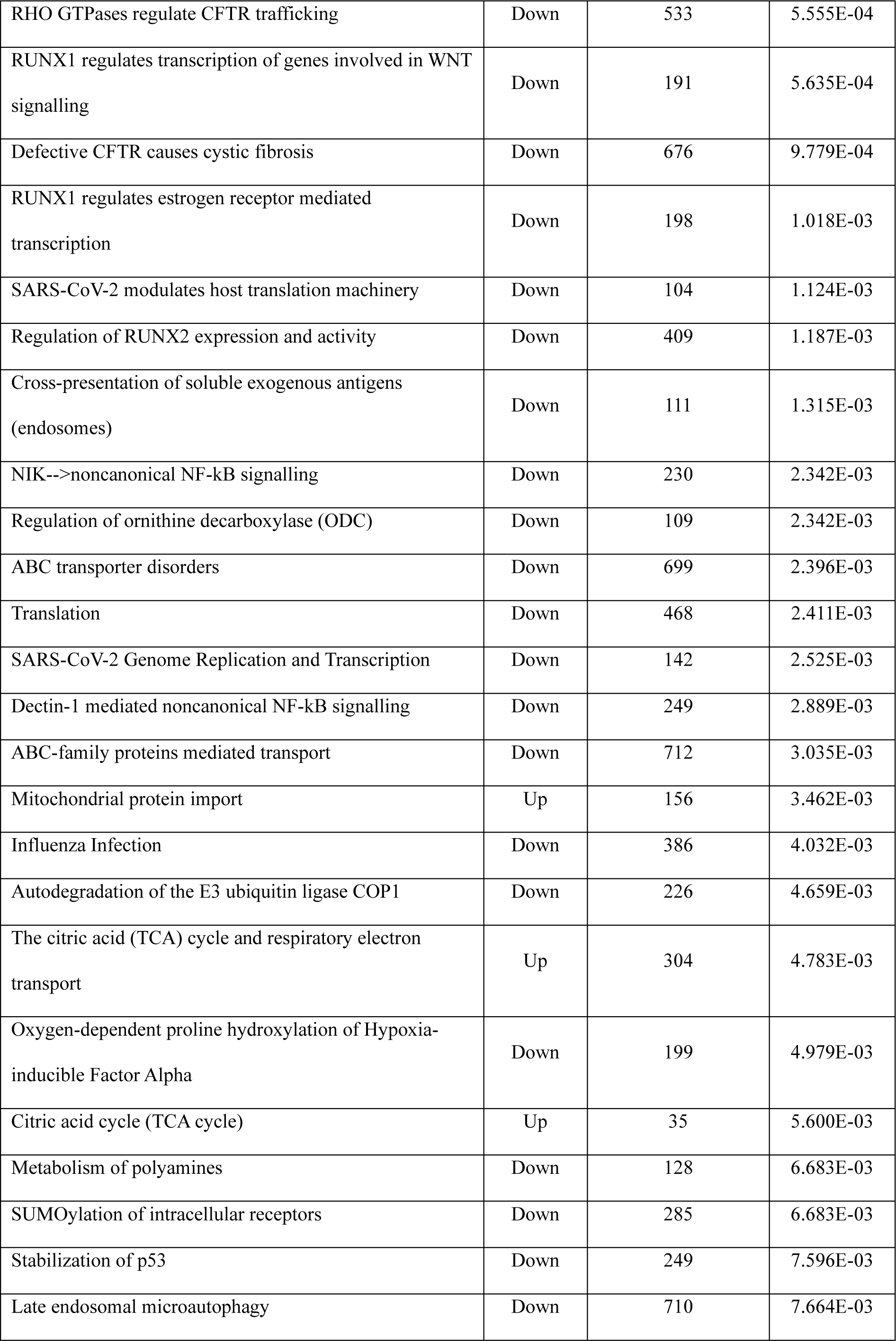
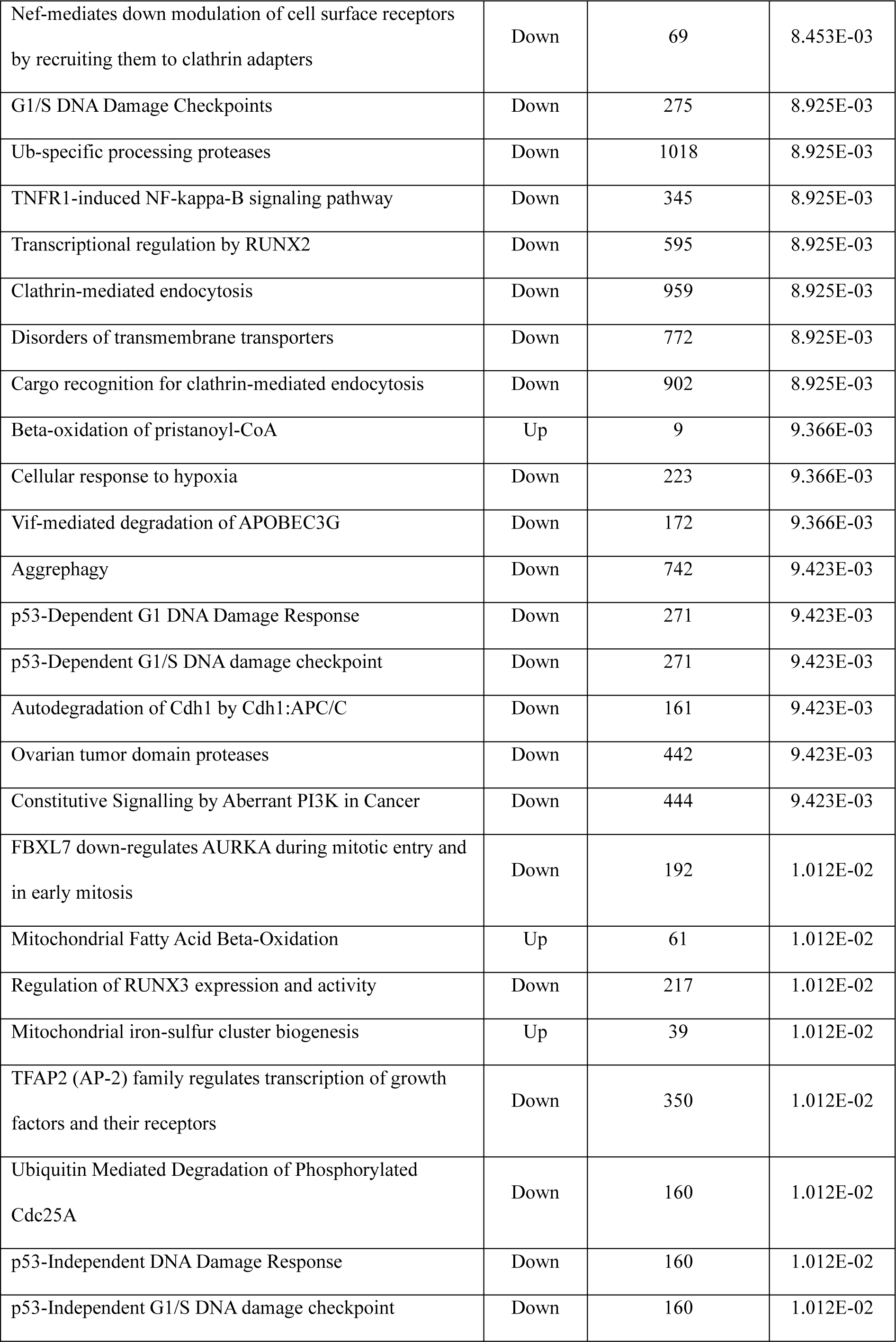
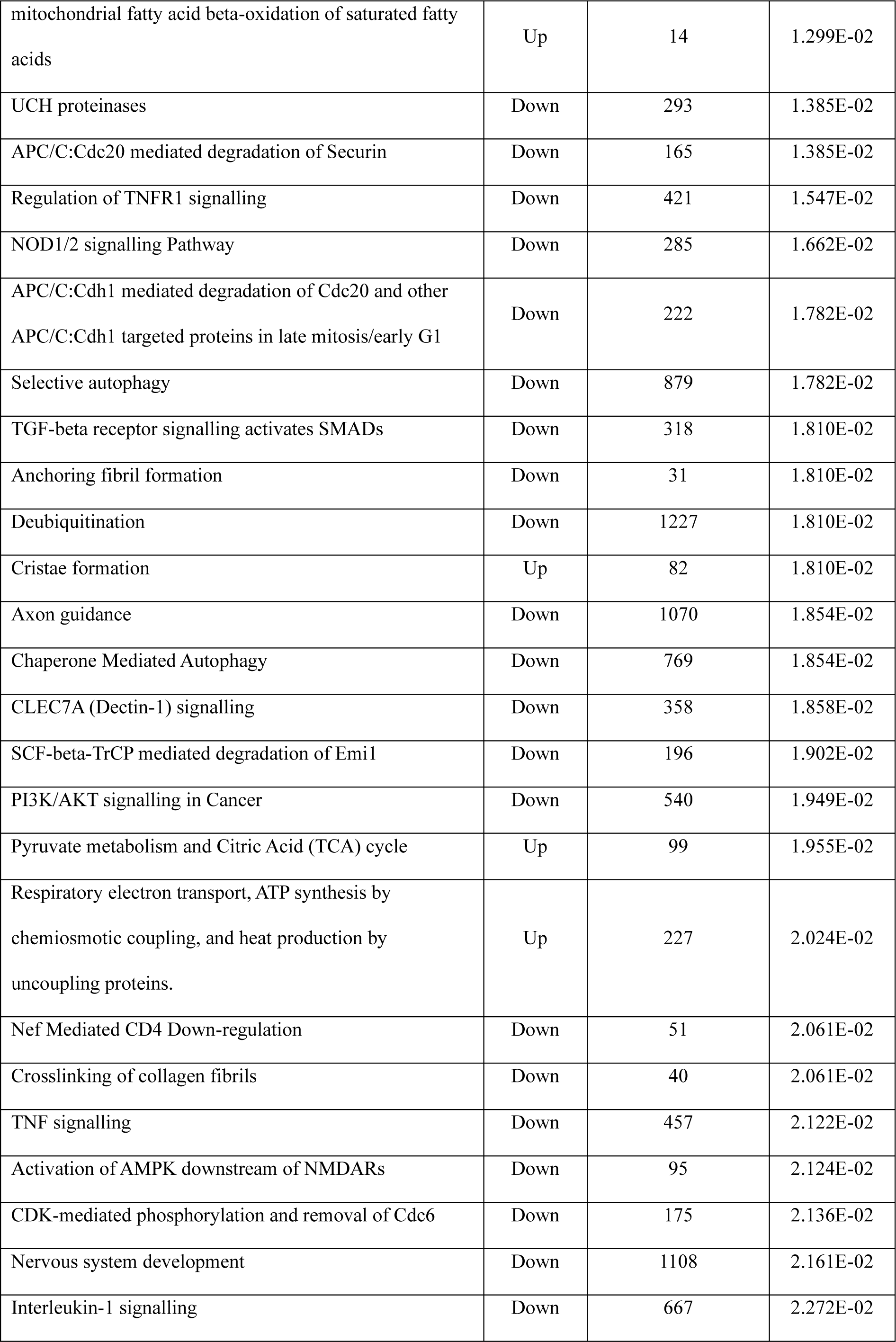
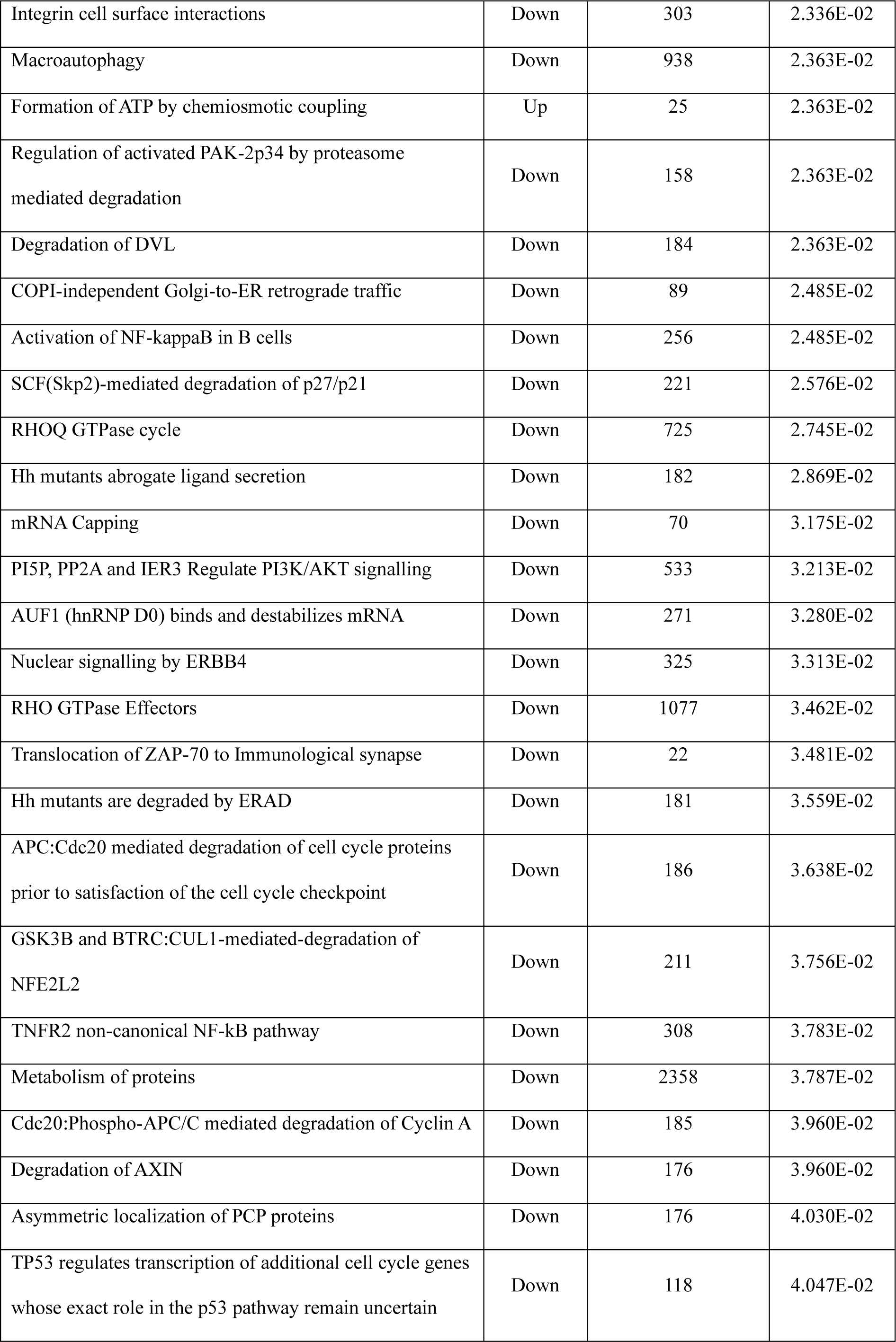
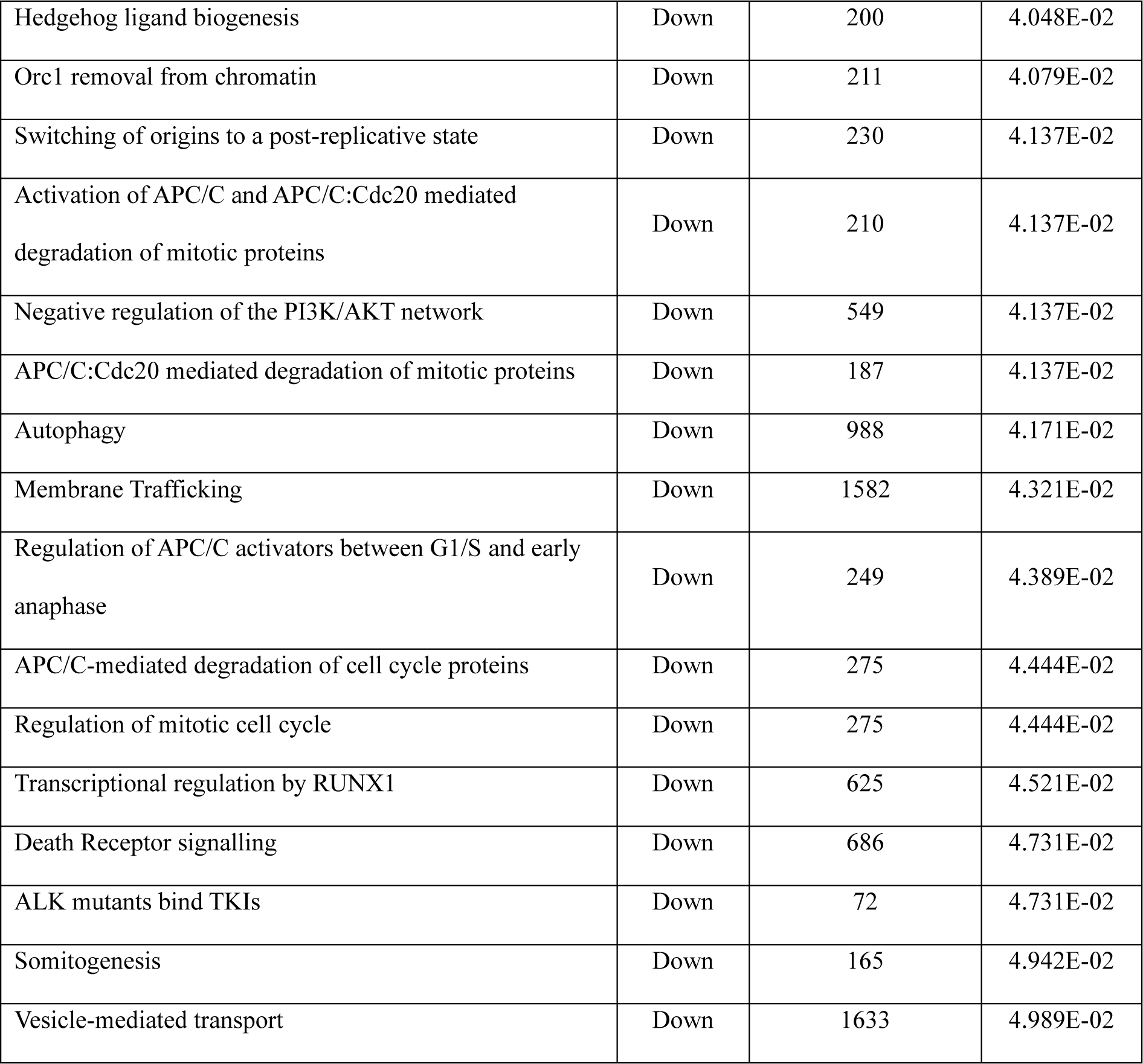
All significantly up- and downregulated Reactome pathways.

